# Tumor neoantigen heterogeneity impacts bystander immune inhibition of pancreatic cancer growth

**DOI:** 10.1101/2020.05.07.083352

**Authors:** Manisit Das, Xuefei Zhou, Yun Liu, Anirban Das, Benjamin G. Vincent, Jingjing Li, Rihe Liu, Leaf Huang

## Abstract

The threshold for immunogenic clonal fraction in a heterogeneous solid tumor required to induce effective bystander killing of non-immunogenic subclones is unknown. Pancreatic cancer poses crucial challenges for immune therapeutic interventions due to low mutational burden and consequent lack of neoantigens. Here, we designed a model to incorporate artificial neoantigens into genes of interest in cancer cells and to test the potential of said antigens to actuate bystander killing. By precisely controlling the abundance of a neoantigen in the tumor, we studied the impact of neoantigen frequency on immune response and immune escape. Our results showed that a single, strong, widely expressed neoantigen could lead to a robust antitumor response when at least 80% of cancer cells express the neoantigen. Further, immunological assays revealed induction of T-cell responses against a non-target self-antigen on KRAS oncoprotein, when we inoculated animals with a high frequency of tumor cells expressing a test neoantigen. Using nanoparticle-based gene therapy, we successfully altered the tumor microenvironment by perturbing interleukin-12 and interleukin-10 gene expression. The subsequent remodeling of the microenvironment reduced the threshold of neoantigen frequency at which bioluminescent signal intensity for tumor burden decreased 1.5-logfold, marking a robust tumor growth inhibition, from 83% to as low as 29%. Our results thus suggest that bystander killing is rather inefficient in immunologically cold tumors like pancreatic cancer and requires an extremely high abundance of neoantigens. However, the bystander killing mediated antitumor response can be rescued, when supported by adjuvant immune therapy.

## Introduction

Immunotherapy is a significant advance in anticancer drug development distinct from conventional chemotherapy and targeted therapy approaches. The antitumor immune response can sustain for a long time after treatment completion and lead to development of long-lived immune memory, allowing prolonged immune activity ^1-3^. Further, immune responses targeted against specific antigens can be expanded to include other antigenic targets, in a phenomenon known as ‘epitope spreading’ (also known as antigen spreading or determinant spreading) ^4-6^.

In recent years, personalized cancer vaccines targeting neoantigens in patients with advanced metastatic disease have been promising in early clinical trials ^7-9^. Vaccine approaches in cancer therapy rely on mounting immune response against mutated proteins unique to cancer cells, protein products of genes selectively expressed in tumor cells, or overexpressed self-antigens. Cancer cells present tumor-associated antigens (TAAs), which are short peptides, at the cellular surface via major histocompatibility complex (MHC) molecules. Neoantigens, commonly understood to be variant peptides derived from mutated proteins, can elicit a robust T-cell immune response due to the absence of thymic elimination of pre-existing T cells against non-self-antigens. Cancers with a high mutational burden such as melanomas, with an average of over 500 coding mutations ^10^, can elicit a higher number of TAA-specific T cells and tumor-infiltrating lymphocytes (TILs). The access to a vast repertoire of mutations can be leveraged in designing potent immunotherapies.

Accumulating evidence indicates the success of neoantigen vaccines relies on efficient epitope spreading. The exact percentage of cells bearing T cells specific for any given neoantigen is a fraction of the total T cell repertoire, which is why we often see cancer vaccines are designed to target multiple neoantigens to be effective before immune response diversification by epitope spreading^4,11,12^. However, previous works have also shown that immune system is capable of selective recognizing and depleting immunogenic subclones in a predominantly non-immunogenic tumor^13^.

There is limited evidence connecting bystander killing of heterogeneous tumor clones to the ‘frequency of the neoantigen’, which we define in the scope of this study as the proportion of the tumor cells that express a specific immunogenic neoantigen. In other words, we do not know the abundance of neoantigen expressing tumor cells required to see bystander killing, translating into a robust antitumor immune response. This can be an important design consideration in neoantigen-based cancer vaccine development. Therefore, we set up a preclinical model to answer this question.

For our study, we chose the orthotopic syngeneic allograft KPC model of pancreatic cancer ^14,15^. KPC is a clinically relevant, genetically engineered mice model (GEMM) for pancreatic ductal adenocarcinoma (PDAC) which is the dominant subtype of pancreatic cancer, with oncogenic driver mutations in KRAS (G12D) and p53 (R172H). PDAC tumors in both humans and KPC mice manifest intricate networks of immunosuppressive tumor-infiltrating leukocytes, and dense desmoplastic stroma, setting steep challenges for most therapeutic interventions. Further, the KPC model has been shown to lack neoantigens, making it a suitable model to incorporate and study response to induced neoantigens, against a background of less immunogenic epitopes ^16^.

Developing neoantigen vaccines in pancreatic cancer remains challenging. Targeting driver mutations like KRAS^G12D^ is appealing, as these mutations are oncogenic and abundantly expressed by cancer cells^17^. Unfortunately, not all driver mutations will translate into strong neoantigens^18^. There are only a few variants of human leukocyte antigen (HLA) molecules (MHC in humans) that will bind and present the KRAS^G12D^ peptide and translate into KRAS-targeted immunotherapy. Since the combination of these HLA variants and the KRAS^G12D^ mutation are expressed by a fraction of the population at large, potential beneficiaries of such therapeutic approaches are also few.

We wanted to test if strong neoantigen initiated immunity can broaden to include nontargeted self-antigens, actuate bystander killing, and lead to tumor regression in pancreatic cancer. We predicted peptides spanning single point mutations on KRAS that are potential neoantigens, using machine learning algorithms. We picked KRAS as a driver gene that is widely expressed on the cancer cells, and we hypothesized that incorporating a neoantigen in KRAS would have a high probability of inducing bystander killing. Our work is important as it answers the question if the frequency of a neoantigen is critical to elicit a robust tumor antitumor response, and if exposure to immunogenic variant peptide can break tolerance to self-antigens or induce bystander killing of non-immunogenic subclones in the tumor.

## Results

### Predicting neoantigens derived from single point mutations on KRAS *in silico*

We aimed to predict strong neoantigens based on mutations of the KRAS oncoprotein. We wrote Python code to construct an *in silico* library of KRAS sequences with single point mutations (**Supplementary Material 1**). This exercise allowed us to generate a library of 3760 [188 (Length of KRAS protein sequence) x 20 (Number of amino acid single letter codes)] unique sequences. We used this library to predict which of these mutations will translate to ‘strong neoantigens’, using machine learning algorithms. See prediction of neoantigens section in materials and methods for more details.

Strong neoantigens are peptides that presented a percentage rank of predicted affinity to MHC-I below 0.5, when compared to a set of 4 x 10^5^ random natural peptides in the database of NetMHC 4.0 server (http://www.cbs.dtu.dk/services/NetMHC/). Affinity threshold for strong binding peptides was 50.00 nM, and for weak binding peptides was 500.00 nM. We selected candidate binders based on %Rank rather than nM Affinity, following the recommendation of the server developers. For H-2K^b^, D153S variant peptide VSDAFYTL had an affinity of 19.34 nM and % rank of 0.06, while D153M variant peptide VMDAFYTL had an affinity of 71.51 nM and a % rank of 0.20. The self-variant peptide VDDAFYTL had a % rank of 7.50, and an affinity of 5598.20 nM for H-2K^b^ (**Fig. 1D**).

**Fig. 1.**
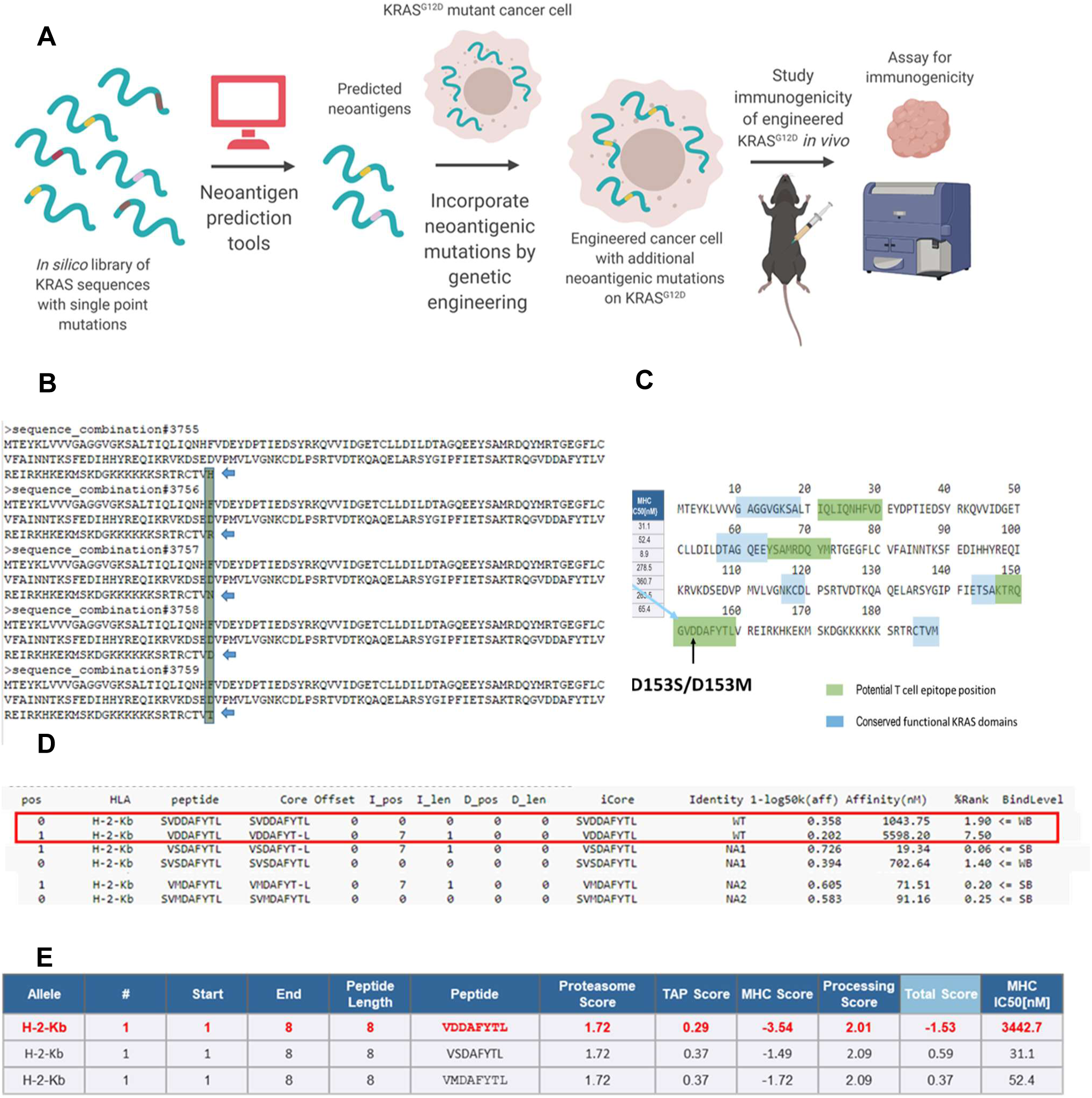
*In silico* prediction of neoantigens derivable from single point mutations on KRAS. a) Scheme of insertion of neoantigens via gene editing, b) Representative sequences from a library of mouse KRAS (UniProt P32883-2) sequences with single amino acid variations, c) Alignment of predicted mutation-derived neoantigens against conserved KRAS functional domains, d) Predicted changes in sequence scoring upon single point mutation, from weak binding level (WB) to strong binding level (SB), via NetMHC 4.0 Server, Technical University of Denmark, http://www.cbs.dtu.dk/services/NetMHC/, e) Predicted changes in sequence scoring upon single point mutation, via Immune Epitope Database and Analysis Resource, http://tools.iedb.org/main/tcell/

We generated a cell line from the genetically engineered KPC mice model^19-21^ with KRAS^G12D^ and p53^R172H^ oncogenic driver mutations, and inoculated cells orthotopically in mice sharing the same genetic background, for two reasons: Firstly, having a non-spontaneous model allowed us to precisely control the frequency of neoantigen to assay the subsequent effect on tumor elimination while reasonably replicating the desmoplastic and immunosuppressive characters of the GEMM tumors. Secondly, it allowed us to engineer the cell line to express luciferase and quantitatively monitor tumor growth by bioluminescence imaging. We tested the effects on tumor growth and immunity in orthotopic animal models. We used nanoparticle-based drug delivery to locally and transiently express immunomodulatory cytokines in the PDAC tumors ^20,21^, and as a tool to probe the effect of TME alterations on bystander killing. A simplified scheme of the approach is presented in **Fig. 1A**. Representative sequences from the library of KRAS oncoprotein with single point mutations are presented in **Fig. 1B**.

We used open-access resources based on artificial neural networks ^22,23^ to predict T cell epitopes restricted to major histocompatibility complex class I (MHC-I) molecules (H-2K^b^, H-2D^b^) expressed in C57BL/6 mice-a strain syngeneic to KPC cell lines. These tools predicted IC50 values for peptide binding to specific MHC molecules, proteasomal processing, Transporter associated with antigen processing (TAP) transport, and produced overall scores indicating the intrinsic potential of a peptide to be a T cell epitope. The epitopes were ranked by potential, and the epitopes overlapping with functional KRAS domains (**Fig. 1C**) were eliminated ^24^, to avoid compromising the functionality of the oncoprotein while changing the immunogenicity. After these exercises, two mutations D153S and D153M were found to have the highest scores to be potential T cell epitopes presented by MHC-I on C57BL/6 mice. The scoring comparison of these epitopes against the wild type sequence were presented for several parameters governing the potential of MHC-I restricted antigen recognition, as listed above (**Fig. 1D-E**). We noted the D153S and D153M mutations significantly improved the scorings for H-2K^b^ restriction, but not for H-2D^b^ restriction (**Supplementary Table 1)**.

### Engineering plasmids to express KRAS oncoprotein with point mutations predicted as neoantigens

We engineered plasmids encoding KRAS oncoproteins with mutations predicted to be MHC-I restricted T cell epitopes (KRAS^G12D, D153S,^ and KRAS^G12D, D153M^) in mice sharing the same genetic background with the KPC model. A scheme is presented in **Fig. 2A**. The validation of the substitutions on nucleotide and amino acid sequences by site-directed mutagenesis in the KRAS open reading frame (ORF), for the gene insert, and the plasmids, are shown in **Fig. 2B, & 2C**. We then chose three different cell lines – a breast cancer cell 4T1 with wild-type KRAS ^25^, colon cancer cell line CT26 with KRAS^G12D 26^, and pancreatic cancer cell line KPC, also bearing KRAS^G12D^ to test the functionality of the mutated KRAS oncoprotein. The results are presented in **Fig. 2D** where we transfected the mutant KRAS^G12D, D153S^ into the 4T1 cells and assayed for p44/42 MAPK (or ERK1/2) protein expression. ERK1/2 is downstream of RAS and known to be activated by mutation in RAS proteins ^27^. We observed transfection with plasmids expressing either KRAS^G12D, D153S^ or KRAS^G12D^ significantly increased ERK1/2 expression relative to transfection with empty vector control, in 4T1 cells. In KRAS-mutant CT26 and KPC cells, we observed no significant difference between untreated cells and cells treated with empty vector or KRAS^G12D^ expressing plasmid. However, transfection with plasmids encoding KRAS^G12D, D153S^ significantly increased ERK1/2 levels over an empty vector or untreated cells, but not over cells treated with plasmids encoding KRAS^G12D^.

**Fig. 2.**
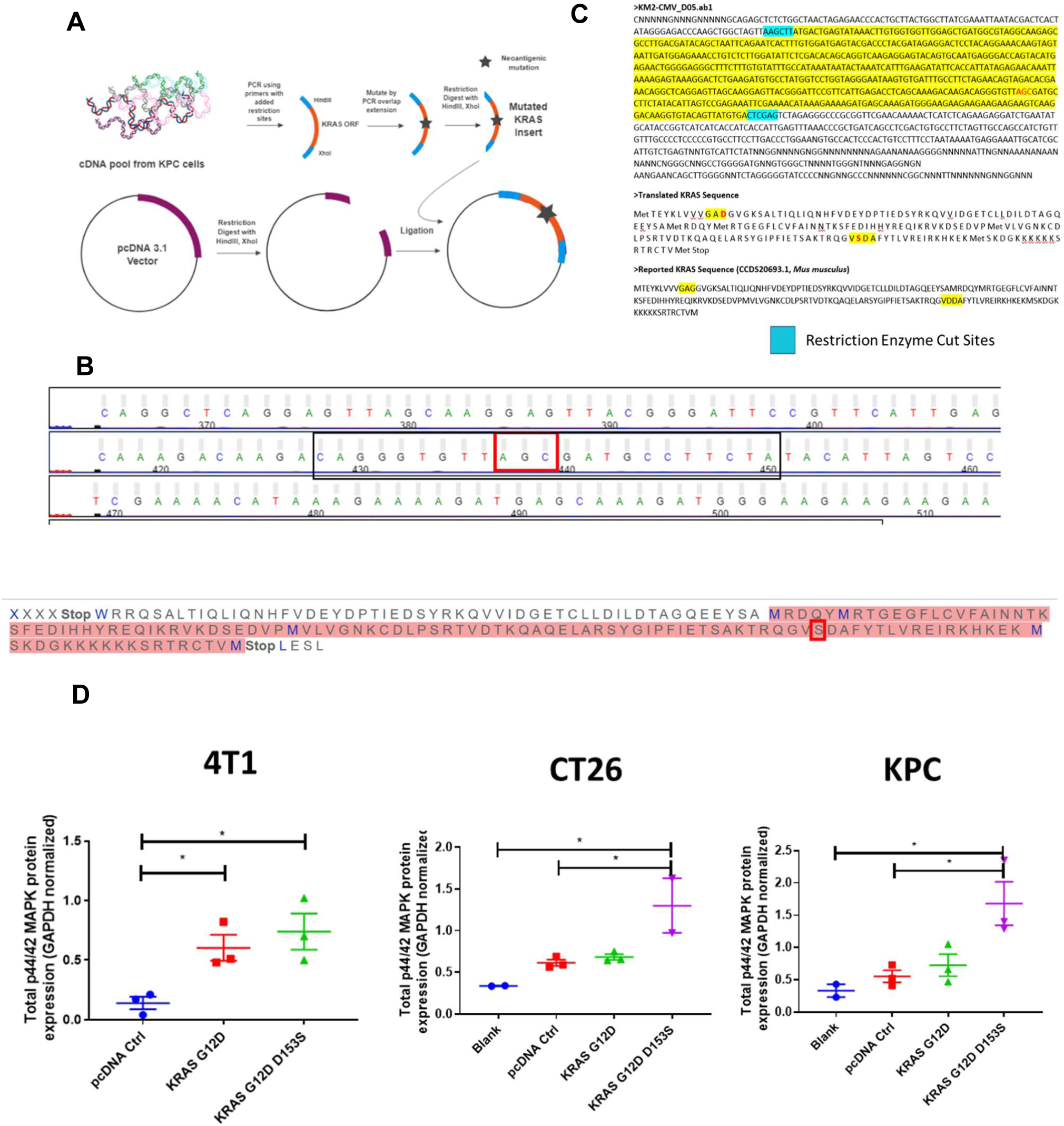
Designing plasmids encoding mutant KRAS and determining functionality of mutant KRAS. a) Workflow for designing plasmids encoding KRAS with mutations predicted to generate neoantigens, mutations were incorporated into DNA inserts via PCR overlap extension, b) Sanger sequencing results confirm incorporation of the D153S mutation-the mutation predicted to have highest binding to MHC-I alleles for C57BL/6 mice, sharing genetic background with KPC, c) Sanger sequencing results confirm incorporation of the D153S mutation in plasmids with pcDNA3.1 backbone, d) Western blot analysis of lysates from *in vitro* cell cultures transfected with KRAS ORF plasmids for 48 h, using Lipofectamine 2000 transfection reagent, analyzing for p44/42 MAPK, downstream of RAS (n= 2-3). Data show mean ± SEM. * p<0.05.

### Investigating the role of neoantigen expression in KPC tumor elimination

We generated monoclonal cell lines expressing KRAS^G12D, D153S^ (cell line henceforth denoted as 3F11) and KRAS^G12D, D153M^ (henceforth as 4E1) respectively. The heterozygous expression of two variants of the KRAS gene in these two cell lines is presented in **Fig. 3A** and **Fig. 3C**. Subsequently, we inoculated KPC cells orthotopically containing a combination of KPC cells expressing wild-type KRAS^G12D^ (denoted henceforth as KPCF1) and either 3F11 or 4E1 respectively, allowed tumors to grow in the pancreas, and monitored tumor growth by bioluminescence imaging. The log-normalized quantification of the radiance, as a measure of tumor growth are presented for mixed inoculates of KPCF1:3F11 in **Fig. 3B**, and for KPCF1:4E1 in **Fig. 3D**. For both combinations with cells expressing D153S or D153M respectively in varying proportions, tumor burden was observed to decrease with an increase in the percentage of cells harboring mutations predicted to be neoantigens, at ten days post tumor inoculation. The log-normalized tumor burden reduced from 7.78 ± 0.10 for tumors with 100% of KPCF1 cells to 6.13 ± 0.16 for tumors with 100% of 3F11 cells (p < 0.0001) (**Fig. 3B**). Similarly, we saw a decrease of log-normalized tumor burden from 7.79 ± 0.11 for tumors with 100% of KPCF1 cells to 6.02 ± 0.37 for tumors with 100% of 4E1 cells (p < 0.001) (**Fig. 3D**).

**Fig. 3.**
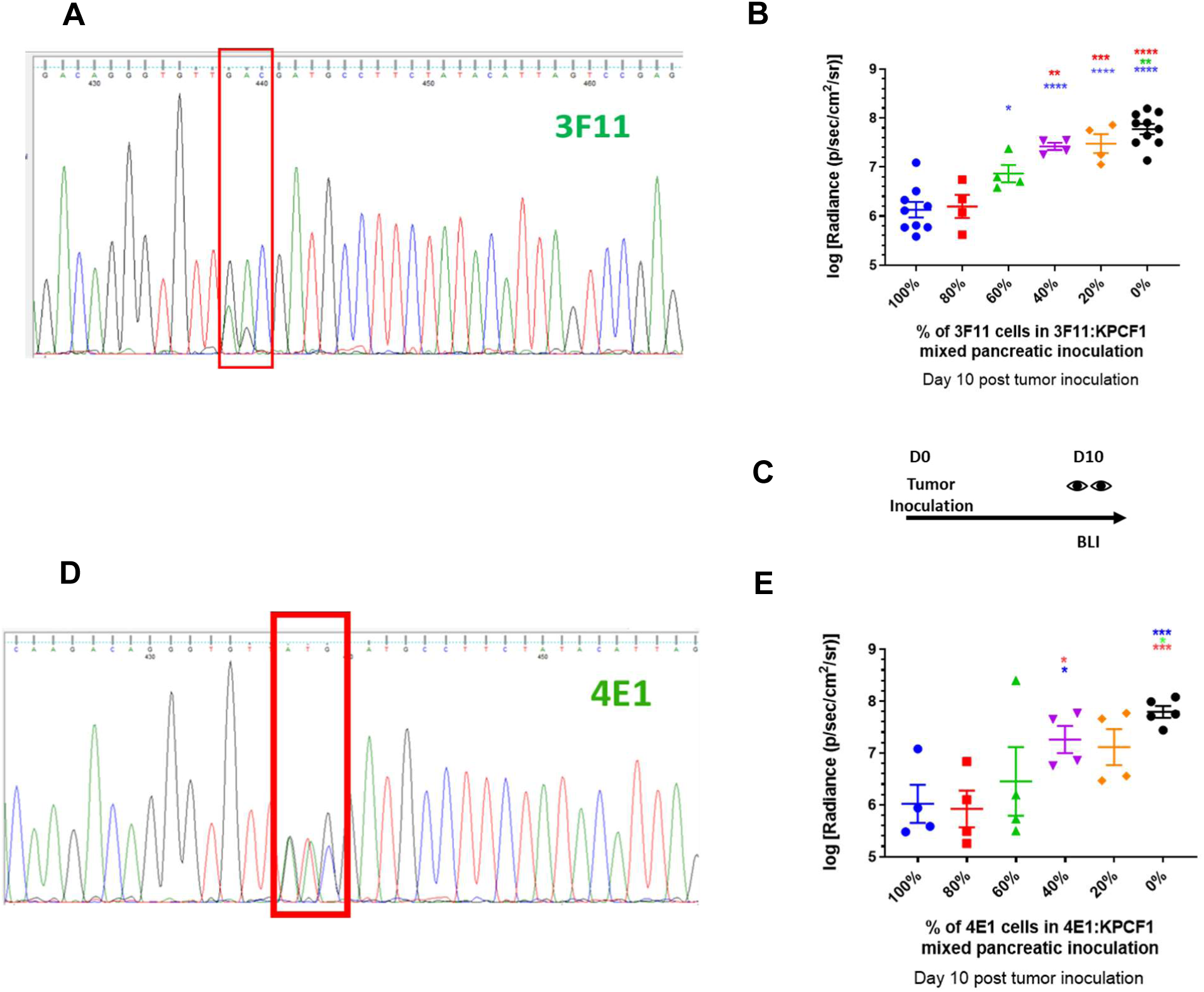
Engineering monoclonal cell lines expressing mutant KRAS and measuring change in antitumor response with variation in epitope frequency. a) Sanger sequencing results confirm expression of the D153S mutation alongside WT allele – in a monoclonal cell line to be referred as 3F11, b) Effect of mutation on orthotopic pancreatic tumor growth via bioluminescence imaging (n = 4-10). Day 10 post tumor inoculation, KPCF1 is a KPC cell line with KRAS^G12D^ mutation, 3F11 is a KPC cell line with KRAS^G12D/D153S^ mutation, c) Imaging protocol for Fig. 3b and 3e. BLI stands for Bioluminescence imaging, and D stands for Day, d) Sanger sequencing results confirm expression of the D153M mutation alongside WT allele – in a monoclonal cell line to be referred as 4E1, e) Effect of mutation on orthotopic pancreatic tumor growth via bioluminescence imaging (n = 4-5). Day 10 post tumor inoculation, KPCF1 is a KPC cell line with KRAS^G12D^ mutation, 4E1 is a KPC cell line with KRAS^G12D/D153M^ mutation. Data show mean ± SEM. * p<0.05, ** p<0.01, *** p< 0.001, **** p<0.0001.

### Evaluating the role of immunotherapy to reduce the threshold of selective outgrowth of antigen-negative tumors

We repeated the experiment presented in **Fig. 3B**, with an additional variable, where all tumors irrespective of their composition were treated with a plasmid (IL-10t) that encodes an ‘IL-10 trap’, an engineered protein capable of blocking interleukin-10 in the tumor ^21^. Here, we delivered the IL-10 trap gene using lipid-protamine-DNA (LPD) nanoparticles (**Fig. 4A**). The treatment schedule is shown on **Fig. 4B**. The significant reduction (p < 0.05) in IL-10 gene expression in KPC tumors with IL-10t treatment was validated by quantitative PCR (**Fig. 4C**). A significant reduction (p < 0.01) in tumor burden was observed for animals harboring tumors with 100% of KPCF1 cells and treated with IL-10t (7.77 ± 0.10) vs. animals treated with phosphate-buffered saline (PBS) (7.20 ± 0.14), however, the reduction was higher for animals harboring tumors with 100% of 3F11 cells, (6.13 ± 0.16 for treatment with IL-10t vs. animals treated with PBS (5.30 ± 0.06) (p < 0.01). A similar trend was observed when tumor-infiltrating immune cells were assayed using flow cytometry. A significant increase of CD3^+^ T cells, CD8^+^ CD3^+^ cytotoxic T lymphocytes (CTLs), CD206^-^ F4/80^+^ M1 macrophages, and CD11c^+^ MHC-II^+^ dendritic cells (DCs) were observed for tumors with 100% of 3F11 cells, under IL-10t treatment **(Fig. 4D)**.

**Fig. 4.**
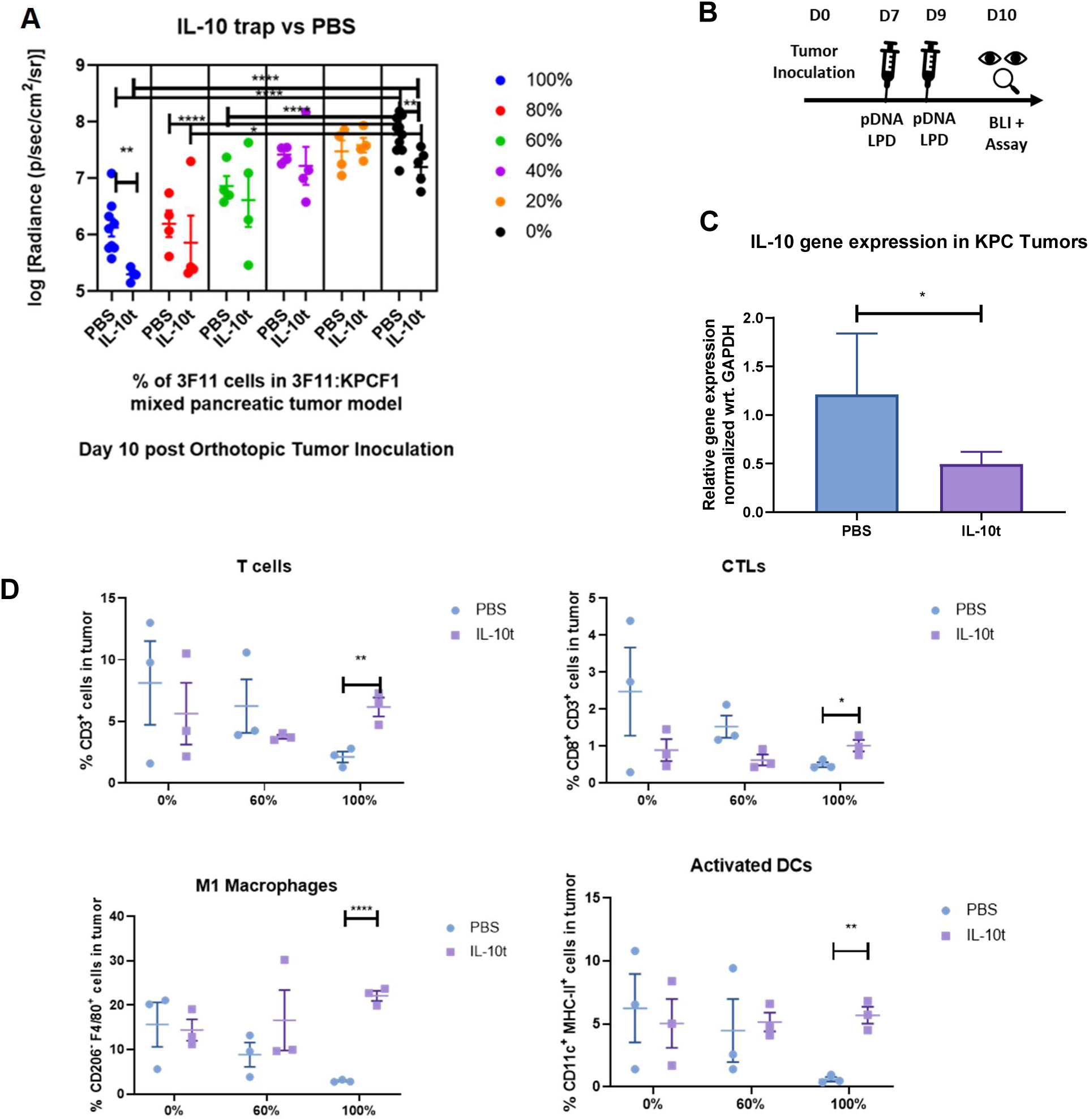
Antitumor response with variation in epitope frequency under local Interleukin 10 blockade. a) Effect of mutation on orthotopic pancreatic tumor growth via bioluminescence imaging (n = 4-10). Day 10 post tumor inoculation, KPCF1 is a KPC cell line with KRAS^G12D^ mutation, 3F11 is a KPC cell line with KRAS^G12D/D153S^ mutation. Animals received either phosphate buffered saline (denoted as PBS) or 50 μg of IL-10 trap plasmid DNA (denoted as IL-10t) administered via lipid-protamine-DNA (LPD) nanoparticles intravenously on Day 7 and 9 post orthotopic tumor cell inoculation, b) Treatment regimen for Fig. 4a. pDNA stands for plasmid DNA and BLI stands for bioluminescence imaging, c) Animals were sacrificed on Day 10 post orthotopic tumor inoculation, tumors were harvested, and mRNA Gene expression were quantified by qPCR (n = 16-18), d) Animals were sacrificed on Day 10 post orthotopic tumor inoculation, tumors were harvested, and immune cells were characterized and quantified by flow cytometry (n = 3). Data show mean ± SEM. * p<0.05, ** p<0.01, *** p< 0.001, **** p<0.0001.

To evaluate potential differences in tumor growth with mixed populations in the context of increased immune activation, we took a similar approach and repeated the tumor growth experiments with varying combinations of KPCF1 and 3F11 cells treated with IL-12 plasmid DNA delivered by LPD nanoparticles or PBS. The quantification of tumor burden during sacrifice at ten days post tumor inoculation is presented in **Fig. 5A**, and the treatment regimen on **Fig. 5B**. We saw a similar trend as we observed in **Fig. 4A**. However, unlike treatment with IL-10t, where we saw a significant alteration in tumor burden only with 100% and 0% of 3F11 cells, here, under IL12 treatment, we observed a significant decrease of tumor burden at 100%, 60%, 40%, 20%, and 0% levels of tumors with 3F11 cells, complemented with KPCF1 cells. We observed over a thousand-fold increase (p < 0.0001) in IL-12 gene expression within the KPC tumors when the gene was delivered using LPD nanoparticles (**Fig. 5C**). We observed a trend in the increase of antitumor immune cells with IL-12 treatment, however, the differences were non-significant (**Fig. 5D**) in the current experimental design (n=3).

**Fig. 5.**
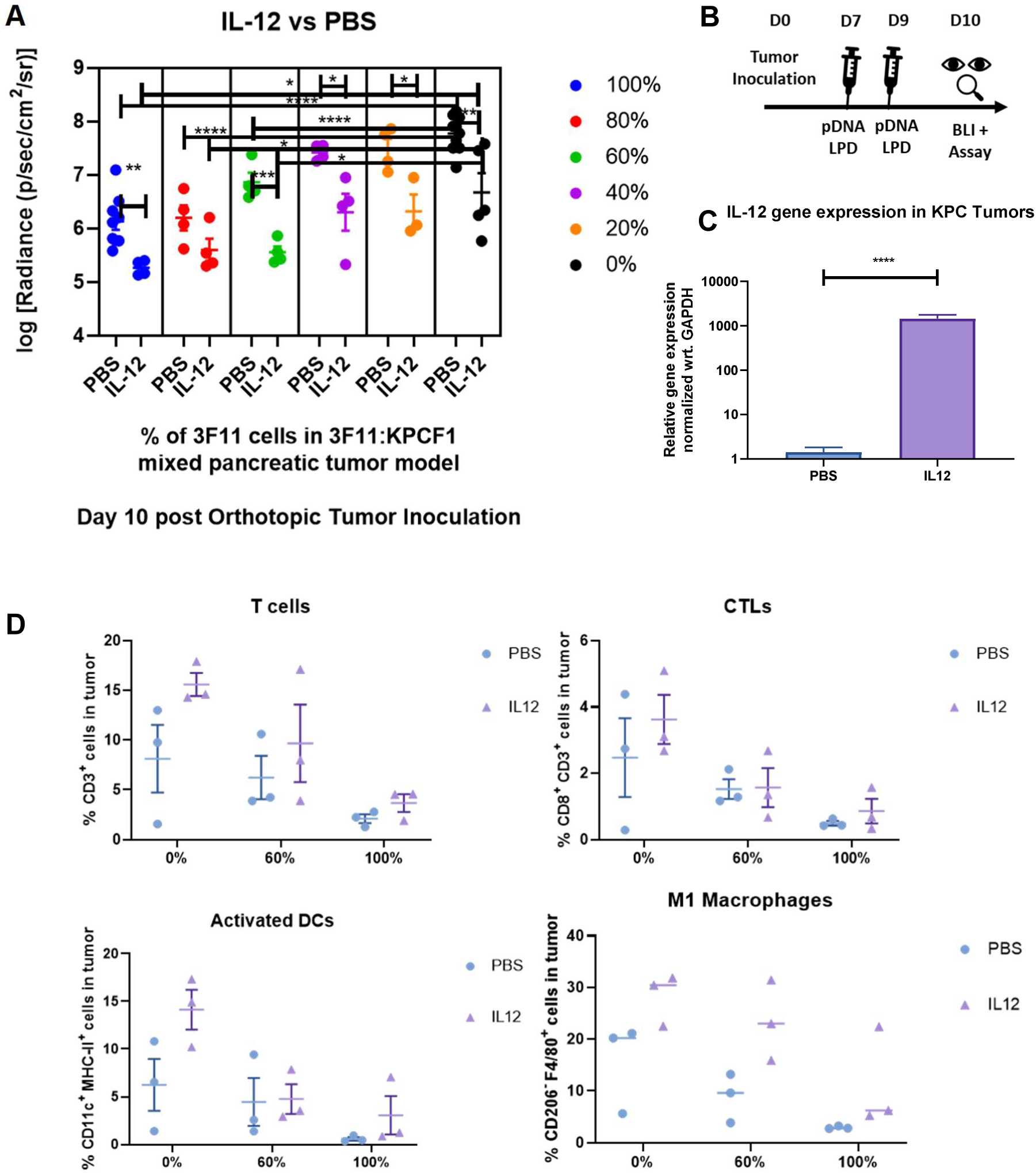
Antitumor response with variation in epitope frequency under local Interleukin 12 gene expression. a) Effect of mutation on orthotopic pancreatic tumor growth via bioluminescence imaging (n = 4-10). Day 10 post tumor inoculation, KPCF1 is a KPC cell line with KRAS^G12D^ mutation, 3F11 is a KPC cell line with KRAS^G12D/D153S^ mutation. Animals received either phosphate buffered saline (denoted as PBS) or 50 μg of IL-12 plasmid DNA (denoted as IL12) administered via lipid-protamine-DNA (LPD) nanoparticles intravenously on Day 7 and 9 post orthotopic tumor cell inoculation, b) Treatment regimen for Fig. 5a. pDNA stands for plasmid DNA and BLI stands for bioluminescence imaging, c) Animals were sacrificed on Day 10 post orthotopic tumor inoculation, tumors were harvested, and mRNA Gene expression were quantified by qPCR (n = 16-18), d) Animals were sacrificed on Day 10 post orthotopic tumor inoculation, tumors were harvested, and immune cells were characterized and quantified by flow cytometry (n = 3). Data show mean ± SEM. * p<0.05, ** p<0.01, *** p< 0.001, **** p<0.0001.

### Breaking tolerance to nontarget self-antigens at high frequency of neoantigens

We used ELISPOT assays to quantify peptide reactive T cells from splenocytes ^28^. We saw a high number of spots representing reactive T cells, when splenocytes from mice inoculated with 100% of 3F11 cells (carrying KRAS^G12D, D153S^) were pulsed with either VSDAFYTL peptide corresponding to the D153S mutation, or the wild type peptide VDDAFYTL. The number of spots were significantly low for splenocytes from mice that were inoculated with a combination of cells with 10% or 0% of 3F11 cells, complemented by KPCF1 cells, compared to mice receiving an inoculation of 100% of 3F11 cells (**Supplementary Fig. 1A**). Representative images of the ELISPOT wells from each tumor type-peptide pulse combination are shown in **Supplementary Fig. 1B**.

### Effective immune adjuvant therapy reduces the threshold of neoantigen frequency required to obtain tumor regression benefit

Finally, to visualize the effect of the predicted T cell epitopes on tumor growth, and under immune intervention, we replotted the results shown in **Fig. 3B, Fig. 4A**, and **Fig. 5A**, together in **Fig. 6.** A third-order polynomial fit was used to generate continuous plots of tumor burden as functions of the percentage of cells carrying predicted neoantigen VSDAFYTL on KRAS^G12D, D153S^. We arbitrarily defined a threshold of 1.5 log-fold reductions in radiance, relative to tumors with no D153S mutation harboring cells and immune intervention (i.e. baseline tumor burden of wild-type tumors composed of 100% KPCF1 cells), to measure the strength of antitumor response under neoantigen presentation. Under no immune intervention, the threshold was reached with 83% of D153S mutation containing cells. Treatment with IL-10t or IL-12 reduced that threshold, to 68%, and 29% respectively.

**Fig. 6.**
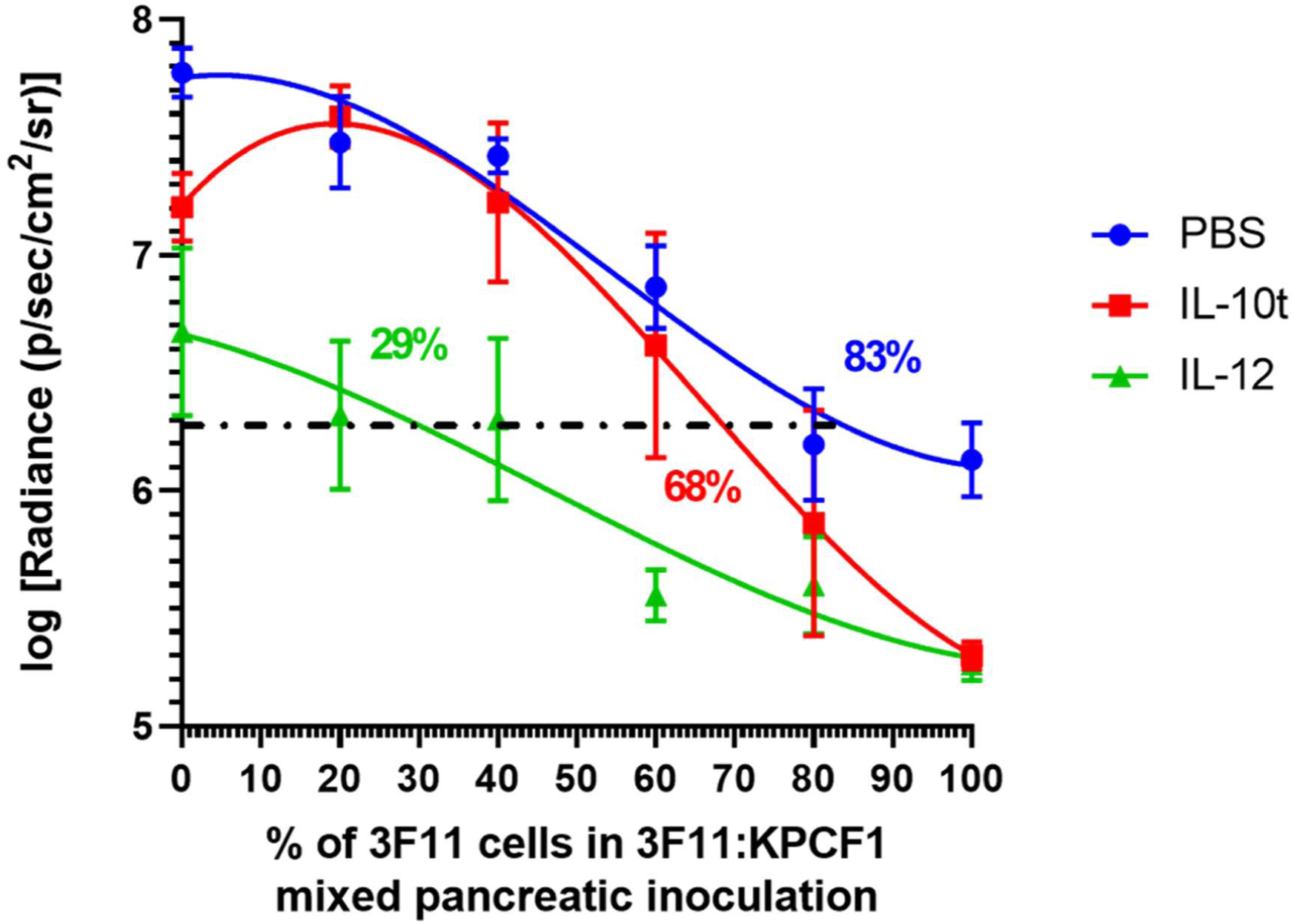
Effective immune adjuvant therapy reduces the threshold of neoantigen frequency required to obtain tumor regression benefit. Effect of mutation and immune intervention on orthotopic pancreatic tumor growth via bioluminescence imaging (n = 4-10). Mice were imaged and tumor signals were quantified on Day 10 post tumor inoculation, KPCF1 is a KPC cell line with KRAS^G12D^ mutation, 3F11 is a KPC cell line with KRAS^G12D/D153S^ mutation. Animals received either phosphate buffered saline (denoted as PBS), 50 μg of IL-12 plasmid DNA (denoted as IL12), or 50 μg of IL-10 trap plasmid DNA (denoted as IL-10t), administered via lipid-protamine-DNA (LPD) nanoparticles intravenously on Day 7 and 9 post orthotopic tumor cell inoculation. The log normalized bioluminescence signal representing tumor burden was plotted on y-axis, and the percentage of cells with D153S mutation in the mixed inoculates were plotted on the x axis. Data fitted using third order polynomial (cubic) interpolation via GraphPad Prism 8.1.0. Horizontal line defines the threshold at which signal intensity for tumor burden reduces 1.5-logfold relative to untreated WT tumor. The numbers on the graph indicate the estimate of neoantigen carrying cells required to reduce the tumor burden by 32-fold or 1.5-logfold in terms of bioluminescence intensity-PBS (89%), IL-10t (68%), and IL-12 (29%). Data show mean ± SEM.

## Discussion

In this study, we aimed to explore the principles that guide tumor neoantigen heterogeneity driven bystander killing of non-immunogenic subclones. Most machine learning algorithms can reliably predict the peptide/MHC binding affinity, although limited in their capabilities to model the immunogenicity of the predicted antigen^29^. The prediction is achieved by modeling high-affinity binding of processed mutated peptides to HLA molecules in humans, and MHC in mice. This approach had been leveraged in the first-in-human neoantigen vaccine clinical trials ^7,9^ to predict strong neoantigens from patients’ tumors, and to subsequently vaccinate the patients with a pool of predicted neoantigens. While these algorithm-driven tools can predict the strength of neoantigen, they currently lack the capacity to assess the frequency of that neoantigen required for epitope spreading. Our model attempted to answer if there is a minimum threshold for neoantigens that is required to elicit a strong antitumor response and break tolerance to self-antigens, and further if such a response can be perturbed by effective immune adjuvant therapy.

Our results showed that incorporating single point mutations on KRAS, predicted to be strong neoantigens, could actuate a robust antitumor response. This emphasized that despite immunosuppressive TME, the immune response was functional and capable of retarding tumor growth in the presence of strong antigens. The effect on tumor growth was further proportional to the frequency of neoantigens, and a sufficiently large number of cells were required to express the neoantigen before a prominent antitumor benefit was observed. In our study, and for the neoantigen we probed, we found that the number is about 83% (**Fig. 6**). Our results were consistent with a previous study ^16^, where ovalbumin (ova) was chosen as an ectopic neoantigen. When animals were inoculated with combinations of ova^+^ and ova^-^ cells in KPC model, tumors with ova^+^ cells were rejected in a CD8^+^ T cell-dependent manner, and selective outgrowth of ova^-^ cells were observed even with 90% of ova^+^ cells in the combination tumors. These results suggest that while the strength of tumor antigenicity is critical in determining fit to be a prospective neoantigen, it is also vital to consider the abundance of the neoantigen, as even a small percentage of cells not expressing neoantigens can escape, and outgrow, a phenomenon that is described as immunoediting^30^. A recent study had also shown that the rejection of immunogenic clones within tumors is dependent on the fractional abundance of the subclone, as well as the nature of the antigen^13^.

We further demonstrated that the threshold at which tumor elimination benefit is lost, and immune escape sets in, can be perturbed by effective immunomodulation. Altering the TME, by increasing the expression of proinflammatory cytokines, or blocking immunosuppressive cytokines, can remarkably reduce the abundance of neoantigen desired for a prominent antitumor response.

It is possible that this bystander killing is augmented by epitope spreading. To enhance the diversity of the tumor-reactive immune repertoire, neoantigen vaccines are commonly designed to target multiple epitopes in the tumor. However, epitope spreading could further broaden the anti-tumor immune response. Both preclinical and clinical evidence points to the existence of epitope spreading, where exposure or vaccination with specific antigens expanded T cell population specific for non-targeted antigens, in melanoma, breast cancer, and prostate cancer ^2,31-35^. In a prominent case, a metastatic melanoma patient was vaccinated with melanoma-associated antigen (MAGE) but manifested only a low level of cytotoxic T cell (CTL) response against MAGE. Most of the T cell receptors recovered from the tumor corresponded to non-vaccine TAAs^2,34^. Together, these evidences suggest epitope spreading as a potential mechanism for tumor regression driven by bystander killing in response to cancer vaccines. Our studies showed tolerance breaking against a self-antigen on KRAS, and in future, we want to investigate if T cell responses are expanded against other non-self-antigens as well, to confirm if bystander killing was mediated by epitope spreading in our approach.

We chose IL-10 and IL12 as model cytokines for intervention, for their importance in the immune microenvironment in PDAC. IL-10 is expressed by immunosuppressive TILs such as tumor-associated macrophages (TAMs) and myeloid-derived suppressor cells (MDSCs), and can result in suppression of dendritic cells (DCs) and subsequent antigen presentation ^36-38^. We had previously demonstrated that blockade of IL-10 in the TME of the KPC model could skew the immunological phenotype towards inflammation ^21^. Here, we administered a subtherapeutic regimen to probe if bystander killing would be actuated when the immunosuppression was reduced. Indeed, the frequency of neoantigens at which a robust tumor response was seen reduced from 83% to 68%. While no significant changes were seen in TILs within tumors containing wild-type KPC cells, tumors that were composed of 100% KRAS^G12D, D153S^ bearing cells had a significant increase in activated DCs after treatment with IL-10t.

IL-12 is capable of reprogramming TME towards an antitumor inflammatory phenotype in several cancers including PDAC. It is known to be a potent molecular vaccine adjuvant and enabler of CTL response, although its use has been limited due to systemic toxicity ^39,40^. Using LPD nanoparticles, we successfully delivered the IL-12 DNA locally and transiently within the PDAC tumor and significantly increased IL-12 gene expression. The subsequent impact on neoantigen-mediated tumor elimination was higher with IL-12 than what we observed with IL-10 trap. The threshold for neoantigen frequency at which a strong response was seen further reduced to 29%, from 83% for tumors receiving no adjuvant therapy.

We also showed that when mice were exposed to orthotopic pancreatic tumors expressing the predicted neoantigen D153S (VSDAFYTL) in high abundance (100%), T cell immune response was developed against the corresponding parental epitope VDDAFYTL (**Supplementary Fig. 1**), which was not seen in mice exposed to only parental D153S negative tumors, or tumors expressing a low abundance of the neoantigen (10%). Previous studies ^11^ showed that when mice were inoculated with tumors engineered to express a specific antigen on one flank, while implanted with parental tumors on the other flank, and vaccinated with the said antigen, both tumors were observed to regress substantially. Our data support the plausibility of such abscopal effect.

In summary, we developed a preclinical model to precisely probe the immune response towards artificial neoantigens; how such an immune response can be affected by the abundance of said neoantigen, and TME modulating interventions. We showed that a single, strong neoantigen expressed by a fraction of tumor cells drive an anti-tumor immune response, although such an approach remained susceptible to immunoediting and selective outgrowth of antigen-poor cell populations. This also suggests that bystander killing is rather inefficient at short-term, and when neoantigen abundance or the fractional abundance of the immunogenic subclone is low. Indeed, bystander killing in the tumor mediated by epitope spreading was reported to have a cascading effect; a previous clinical study showed the number of nontarget antigen recognized increased from 52 to 162, between two to ten weeks post-vaccination against target antigens ^4,32^. In our study, we had chosen a relatively shorter time period which may further explain why the efficiency of bystander killing was low. However, when treating patients with advanced, metastatic disease, time is crucial, and a robust immune response early on is critical to impact survival. On that note, we also established that immunomodulatory therapy targeting IL-10 or IL-12 can significantly shift the threshold for bystander killing, and sustain an antitumor immune response, at a significantly lower abundance of neoantigens. In a clinical setting, our findings suggest when the TME is immunosuppressive, or fractional abundance of an immunogenic subclone is low, bystander killing can be rather inefficient, and can only progress when combined with adjuvant immune therapy.

## Materials and Methods

### Materials

N-(Methylpolyoxyethylene oxycarbonyl)-1,2-distearoyl-sn-glycero-3-phosphoethanolamine, sodium salt (DSPE-PEG 2000, PEG chain molecular weight: 2000) and 1,2-dioleoyl-3-trimethylammonium-propane chloride salt (DOTAP) were procured from NOF Corporation (Tokyo, Japan). Cholesterol was sourced from Sigma-Aldrich (St. Louis, MO, USA). Carbenicillin Disodium Salt, Kanamycin Sulfate, and Matrigel matrix for tumor inoculation were obtained from Corning (NY, USA). Primers for PCR overlap extension, and sanger sequencing were custom ordered from Integrated DNA Technologies, Inc. (Coralville, IA, USA). Polymerase for cloning was obtained from Kapa Biosystems (Wilmington, MA, USA). pcDNA™3.1 (+) Mammalian Expression Vector, DH5α competent *E. coli*, and Lipofectamine™ 2000 Transfection Reagent for cloning and transfection were procured from ThermoFisher Scientific (Carlsbad, CA, USA). Geneticin™ Selective Antibiotic (G418 Sulfate) for selection of monoclonal lines expressing gene of interest was also purchased from ThermoFisher Scientific. Restriction enzymes, T4 DNA ligase, and compatible buffers were purchased from New England BioLabs (Ipswich, MA, USA). Antibodies for Western Blot [Phospho-p44/42 MAPK (Erk1/2) (Thr202/Tyr204) Antibody #9101] were procured from Cell Signaling Technology (Danvers, MA, USA). Antibodies for flow cytometry were procured from BioLegend and eBioscience (San Diego, CA, USA). TaqMan® Gene Expression Assays for quantitative PCR were purchased from ThermoFisher Scientific. Murine IL-12 plasmid DNA was procured from Genscript (Piscataway, NJ).

### Statistical Analyses

We reported data as mean ± standard error of mean (SEM), and replicates for each experiment as ‘n’. GraphPad Prism was used for statistical analyses. Student’s t-test was used for comparing two sets of values, and one-way-analysis-of-variance (ANOVA) for three sets of values and above, with Tukey’s multiple comparisons test for pairwise comparisons. Two-way ANOVA with Tukey’s multiple comparisons test was used for more than one categorical independent variable. *, **, ***, and **** designates p < 0.05, 0.01, 0.001, and 0.0001.

### Generation of *in silico* library of possible KRAS sequences with single point mutations

We used a Python code (See Supplementary Material 1) to generate all possible combinations of mutated sequences from a given library of length N (validAlphabet) i.e. a list of permissible letters designating the single letter codes for amino acids (GALMFWKQESPVICYHRNDT), and a candidate sequence representing the wild type sequence of KRAS oncoprotein available from Consensus CDS (CCDS) project of National Center for Biotechnology Information (CCDS20693.1), as a string input.

We intended to select one letter of the candidate sequence at a time, representing each position of the protein sequence, and substitute it with different letters from the validAlphabet to create different mutated sequence strings, each representing a unique variation of the KRAS protein sequence with a single point mutation in the amino acid level. Thus, we use all letters of the validAlphabet in such a manner to create N possible variations for each letter of the candidate sequence string. We then move on to the next letter in the candidate sequence and repeat the same procedure. Once we create all possible unique combinations i.e. mutated sequences by substituting all letters from the validAlphabet with the selected letter in the candidate sequence, we save it in a text file with .fasta extension in an alphabetically sorted order for downstream analysis.

### Prediction of Neoantigens

T cell epitopes were predicted using open source tools based on artificial neural networks ^22,23^ hosted by Immune Epitope Database (IEDB.org) and NetMHC 4.0 Server (http://www.cbs.dtu.dk/services/NetMHC/). MHC haplotypes corresponding to strain C57BL/6 were used for this exercise as the KPC model of pancreatic cancer chosen for this study shares the same genetic background as C57BL/6 mice. All the 3760 sequences which are variations of KRAS oncoprotein sequences with single amino acid mutations, were used as input sequences for prediction of T cell epitopes. Predicted epitopes were ranked, and potential epitopes were aligned against conserved functional domains on KRAS oncoprotein.

### Design of plasmids encoding KRAS with mutations predicted to generate neoantigens

PCR-driven overlap extension technique was used to perform site-directed mutagenesis and encode the single point mutations predicted as strong T cell epitopes against MHC alleles expressed on C57BL/6 mice, on KRAS oncoprotein ^41^. The original protocol in the cited source was followed with some variations in reagents such as the use of KAPA2G Robust PCR Kit. Briefly, overlapping gene segments were generated using internal primers introducing nucleotide substitutions to incorporate mutations predicted to generate neoantigens-D153S (5’-CAGGGTGTT**AGC**GATGCCTTCTA-3’, 5’-TAGAAGGCATCGCTAACACCCTG-3’), and D153M (5’-CAGGGTGTT**ATG**GATGCCTTCTA-3’, 5’-TAGAAGGCATCCATAACACCCTG-3’), using KRAS open reading frame gene fragment as a starting template.

The KRAS open reading frame starting template DNA was amplified from KPC cells harboring G12D mutation. These overlapping strands were then hybridized at their 3’ region in a second round of PCR and extended to generate a full length KRAS oncoprotein sequences with single point mutations (G12D, D153S/D153M). Restriction enzyme sites were included to enable insertion of the KRAS open reading frame product into expression vector for subsequent cloning. The mutated KRAS DNA fragments were subsequently inserted into pcDNA™3.1 (+) mammalian expression vector after the insert and the vector were gel purified, phosphorylated, and ligated downstream of CMV promoter with HindIII and XhoI restriction sites.

### DNA Sequencing

Sanger sequencing was performed at Genewiz-an R&D genomics service provider. Sample submission guidelines by the provider was follower. For KRAS open reading frame PCR products, one of the primers flanking the two ends of the DNA were used (5’-CAGACTAAGCTTATGACTGAGTATAAACTTGTG-3’, 5’-CAGACTCTCGAGTCACATAACTGTACACCTTGTCCTT-3’). For pcDNA plasmids, primer against the CMV promoter was used for sequencing (5’-CGCAAATGGGCGGTAGGCGTG-3’). Trace data from sanger sequencing were viewed and analyzed with the desktop application FinchTV 1.4.0.

### Analysis of protein expression in cell cultures transfected with KRAS ORF plasmids *in vitro*

Plasmids capable of expressing either control KRAS oncoprotein (KRAS^G12D^), or oncoproteins with predicted neoantigens (KRAS^G12D, D153S^ or KRAS^G12D, D153M^) were transfected into 4T1 (ATCC® CRL-2539™), CT26-FL3^42^, and KPC ^19^ cell lines *in vitro* using Lipofectamine 2000. Transfection Reagent manufacturer’s instructions were followed for transfection. After incubation with transfection reagents for 48 h, lysates were collected from cell culture, using RIPA Lysis and Extraction Buffer, and protein concentrations were determined using bicinchoninic acid (BCA) assay. Western blots were performed with denatured, reduced samples, transferred to membranes and stained with antibodies against p44/42 MAPK. The membranes were developed using HRP-conjugated antibodies enhanced chemiluminescence kits, digitally imaged and signals were quantified using ImageJ. GAPDH was used as an endogenous control.

### Expansion of monoclonal cell lines expressing mutant KRAS

Geneticin® (G-418 Sulfate) was used to generate stably transfected cell lines from *in vitro* KPC cell cultures transfected with pcDNA plasmids encoding KRAS open reading frame. After 48 h of incubation with plasmids complexed with Lipofectamine 2000, cells were incubated with fresh complete media supplemented with G-418 sulfate (500 µg/mL). The media was replaced every 3-4 days with fresh media until the selected cells started growing stably and then selection was maintained by culturing cells in presence of the antibiotic. The expression of the mutations predicted to be antigens (D153S and D153M) were confirmed with Sanger sequencing.

### Cell lines

Primary tumor cell lines of pancreatic ductal adenocarcinoma were derived from a genetically engineered mouse model (LSL-Kras G12D/ +; LSL-Trp53R172H/ +; Pdx-1-Cre, syngeneic to C57BL/6 strain), and obtained as a generous gift from Dr. Serguei Kozlov from Center for Advanced Preclinical Research, Frederick National Laboratory for Cancer Research (NCI). Cells were cultured in Dulbecco’s Modified Eagle Medium: Nutrient Mixture F-12 (DMEM/F12) and supplemented with fetal bovine serum (FBS, 10%) (Gibco), Penicillin/Streptomycin (1%) at 37°C and 5% CO2 in a humidified atmosphere. The primary cell lines were stably transfected with lentiviral vector carrying mCherry red fluorescent protein (RFP) and firefly luciferase (Luc). The stably transfected cell lines (KPCF1) were used for *in vivo* studies and monitored by bioluminescence. The KPCF1 cells were further transfected with pcDNA plasmids bearing KRAS open reading frame DNA fragments, resulting in cells expressing double mutations on KRAS- (G12D, D153S) referred as 3F11, and (G12D, D153M) referred as 4E1. Gene expression was maintained using G-418 Sulfate.

4T1 cells were cultured according to the supplier’s instructions (ATCC® CRL-2539™). CT26-FL3 cells were obtained from Dr. Maria Pena at the University of South Carolina ^42^ and cultured in Dulbecco’s Modified Eagle’s Medium (DMEM) containing 4.5 g/L glucose, and further supplemented with 10% FBS and 1% penicillin/streptomycin at 37°C and 5% CO2 in a humidified incubator.

### Orthotopic pancreatic tumor model to study effect of mutation on tumor growth

All animal experiments were conducted in compliance with regulations of the University of North Carolina at Chapel Hill Institutional Animal Care and Use Committee (IACUC). Orthotopic pancreatic tumors were inoculated as previously described ^19^. Sub-Confluent KPCF1, 3F11, and 4E1 cells were trypsinized, washed in ice-cold PBS, and resuspended in 1:1 mixture of Matrigel Matrix (Corning): phosphate buffered saline (PBS). The cells were mixed in various proportions to tune the frequency of cells varying predicted epitopes and injected in the pancreas of 8-10-week-old C57BL/6 mice anesthetized with isoflurane (10^6^ cells per mice in a volume of 50 μL. USP grade Meloxicam (Thomas Scientific) was administered as a post-operative analgesic.

### Bioluminescence imaging to monitor tumor growth

Tumor growth was monitored by bioluminescence imaging using an IVIS Lumina Series III *In vivo* imaging system (PerkinElmer). Anesthetized animals were administered D-luciferin (100 mg/kg of body weight), intraperitoneally and bioluminescence was recorded five minutes past administration. Bioluminescence signal intensity was reported as radiance- a measurement of photons emitted from the subject, in the units of photons/second/cm^2^/steradian. The radiance values were log-normalized and tabulated.

### ELISPOT Assay to measure IFN-γ response

Animals were sacrificed on Day 10 post tumor inoculation, and spleens were harvested. IFN-γ ELISPOT Assay were performed with splenocytes isolated from spleens, using BD ELISPOT (BD Biosciences, San Jose, CA, US) reagents, following manufacturer’s instructions. The plates were dried, and the number of spots were quantified using AID ELISPOT reader (Autoimmun Diagnostika GmbH, Strassberg, Germany). The peptides for pulsing the splenocytes were procured from Peptide 2.0 (Chantilly, VA, US).The peptide-splenocyte incubation was performed for 18 h at 37 °C in a humidified incubator.

### *In vivo* delivery of plasmid gene using lipid-protamine-DNA (LPD) nanoparticles

LPD nanoparticles were synthesized using a sequential self-assembly process based on a method, described previously ^43^. Briefly, solvent from an equimolar solution of DOTAP and cholesterol in methylene chloride was removed under nitrogen flow. The resultant thin lipid film was hydrated with deionized water, vortexed, and sequentially extruded through 400, 200, and 100 nm polycarbonate membranes (Millipore, MA) to generate unilamellar liposomes with a hydrodynamic size between 100-200 nm. The polyplex cores were formed between 50 μg of plasmid DNA and 22 μg of protamine in 5% glucose (plasmids encoding IL-10 trap, as described in our previous work ^21^, or IL-12 that was commercially purchased). The core complex was incubated for ten minutes at room temperature and then added to 60 μL of DOTAP/Cholesterol liposomes (10 mM each). Postinsertion of 15% DSPE-PEG 2000 was performed at 65 °C for 15 min, and the final nanoparticles were administered to animals intravenously.

### Characterization of tumor-infiltrating immune cells using flow cytometry

Post-sacrifice, orthotopic pancreatic tumors were harvested and enzymatically digested with Type IV Collagenase (Gibco) and Deoxyribonuclease I (Alfa Aesar). The samples were subsequently passed through 40 µm cell strainer (Corning), washed with and resuspended in Fluorescence-activated cell sorting (FACS) buffer (PBS supplemented with 10% FBS, and 2 mM EDTA). Cells were stained with fluorescent-conjugated antibodies and fluorescence parameters were recorded with Becton Dickinson LSR II (HTS) flow cytometry analyzer. OneComp eBeads™ Compensation Beads were used for single-color compensation controls. Data analyzed using FlowJo V10.

### Quantification of mRNA gene expression by quantitative PCR

Pancreatic tumor tissues were harvested, and total RNA was isolated using RNeasy kit (Qiagen). Complementary DNA (cDNA) was reverse transcribed using iScript cDNA Synthesis Kit (Bio-Rad). Quantitative polymerase chain reaction (qPCR) was conducted using TaqMan™ Universal Master Mix II, with Uracil-N-glycosylase and TaqMan™ gene expression assays (Applied Biosystems). The following assays were used for amplification of the genes of interest with mouse as target species-GAPDH (Mm99999915_g1), IL-12a (Mm00434165_m1), and IL-10 (Mm01288386_m1). GAPDH was used as endogenous control. PCR reactions were performed with 7500 Real-Time PCR system (Applied Biosystems) and data analyzed with the associated 7500 software.

## Acknowledgments

The work was supported by NIH grant CA198999 and the Eshelman Institute for Innovation. We appreciate the helpful guidance from Dr. Yuhua Wang on experimental designs for cloning experiments. The UNC Flow Cytometry Core Facility is supported in part by P30 CA016086 Cancer Center Core Support Grant to the UNC Lineberger Comprehensive Cancer Center.

## Author Contributions

MD designed and conducted experiments, interpreted data and wrote manuscript; XZ, YL, and JL conducted experiments; AD wrote code; BGV, LH, and RL interpreted data and wrote manuscript.

## Data Availability Statement

The datasets generated during and/or analyzed during the current study are available from the corresponding author on reasonable request.

## Code Availability Statement

The source code generated for this study is provided in the supplementary information.

