## Supplementary Information for "Tumor neoantigen heterogeneity impacts bystander immune inhibition of pancreatic cancer growth"

### Supplementary Material 1. Source Code to generate *in silico* library of KRAS sequences with single point mutations

```
import csv
from textwrap import TextWrapper

def writeToTextFile(combinations, filename):  #if you want text file
    wrapper = TextWrapper(width=80)
    filepath = 'Outputs/' + filename + '.fasta'
    with open(filepath, 'w') as myfile:
        for element in range(len(combinations)):
            myfile.write(">sequence_combination#{}x\n"\
                         .format(element))
            myfile.write("\n".join(wrapper.wrap(combinations[element])))
            myfile.write("\n")
    print("Generated text file :", filename+'.fasta')

#==================================================================================
def writeToCSV(combinations, filename): #if you want csv
    filepath = 'Outputs/'+ filename + '.csv'
    delimeter = "," #enter the delimeter you want
    with open(filepath, 'w') as myfile:
        wr = csv.writer(myfile,delimiter= delimeter, quoting=csv.QUOTE_NONE)
        wr.writerow(combinations)
    print("Generated CSV file :", filename+'.csv')

if __name__ == '__main__':
    validAlphabet = 'GALMFWKQESPVICYHRNDT'
    file = open('sequenceFile.conf', 'r')
    proteinSeq = file.read().strip().split('\n')
    file.close()
    for sequence in proteinSeq:
        sequenceName, actualSequence = sequence.split('=')
        allCombinations = list()
        for i in range(len(actualSequence.strip())):
            for letter in validAlphabet:
                dummySeq = list(actualSequence.strip())
                dummySeq[i] = letter
                dummySeq = ''.join(dummySeq)
                allCombinations.append(dummySeq)
```

```
    uniqueCombinations = list(set(allCombinations)) #unique combinations
    uniqueCombinations = sorted(uniqueCombinations) #sort the set
    totalUniqueGenerated = len(uniqueCombinations)  #number of unique combinations
    print("\nNumber of entries generated for sequence: {} =
{}".format(sequenceName,totalUniqueGenerated))
    name_of_generated_File = 'proteinSequence_'+ sequenceName.strip() + '_'+
str(totalUniqueGenerated)
    #generate text file
    writeToTextFile(uniqueCombinations, name_of_generated_File)
    #generate CSV file
    writeToCSV(uniqueCombinations, name_of_generated_File
```

**Supplementary Table 1. MHC-I Processing Prediction Results**

The MHCI binding predictions were made using the IEDB analysis resource Consensus tool [1] which combines predictions from ANN aka NetMHC (4.0) [2][3][4], SMM [5] and Comblib [6].


1. *Kim Y, Ponomarenko J, Zhu Z, Tamang D, Wang P, Greenbaum J, Lundegaard C, Sette A, Lund O, Bourne PE, Nielsen M, Peters B. 2012. Immune epitope database analysis resource. NAR.*
2. *Nielsen M, Lundegaard C, Worning P, Lauemøller SL, Lamberth K, Buus S, Brunak S, Lund O. 2003. Reliable prediction of T-cell epitopes using neural networks with novel sequence representations. Protein Sci 12:1007-1017.*
3. *Lundegaard C, Lamberth K, Harndahl M, Buus S, Lund O, and Nielsen M. 2008. NetMHC-3.0: Accurate web accessible predictions of Human, Mouse, and Monkey MHC class I affinities for peptides of length 8-11. NAR 36:W509-512.*
4. *Andreatta M. and Nielsen M. 2016. Gapped sequence alignment using artificial neural networks: application to the MHC class I system. Bioinformatics 32:511-7.*
5. *Peters B, Sette A. 2005. Generating quantitative models describing the sequence specificity of biological processes with the stabilized matrix method. BMC Bioinformatics 6:132.*
6. *Sidney J, Assarsson E, Moore C, Ngo S, Pinilla C, Sette A, Peters B. 2008. Quantitative peptide binding motifs for 19 human and mouse MHC class I molecules derived using positional scanning combinatorial peptide libraries. Immunome Res 4:2.*


| **Allele** | **#** | **Start** | **End** | **Peptide Length** | **Peptide** | **Proteasome Score** | **TAP Score** | **MHC Score** | **Processing Score** | **Total Score** | **MHC IC50** |
|---|---|---|---|---|---|---|---|---|---|---|---|
| H-2-Kb | 1 | 1 | 8 | 8 | V**S**DAFYTL | 1.72 | 0.37 | -1.49 | 2.09 | 0.59 | 31.1 |
| H-2-Kb | 2 | 1 | 8 | 8 | V**M**DAFYTL | 1.72 | 0.37 | -1.72 | 2.09 | 0.37 | 52.4 |
| H-2-Kb | 3 | 1 | 8 | 8 | V**D**DAFYTL | 1.72 | 0.29 | -3.54 | 2.01 | -1.53 | 3442.7 |
| H-2-Db | 2 | 1 | 8 | 8 | V**M**DAFYTL | 1.72 | 0.37 | -4.27 | 2.09 | -2.18 | 18786.9 |
| H-2-Db | 1 | 1 | 8 | 8 | V**S**DAFYTL | 1.72 | 0.37 | -4.36 | 2.09 | -2.27 | 22851.5 |
| H-2-Db | 3 | 1 | 8 | 8 | V**D**DAFYTL | 1.72 | 0.29 | -4.57 | 2.01 | -2.56 | 37163.8 |

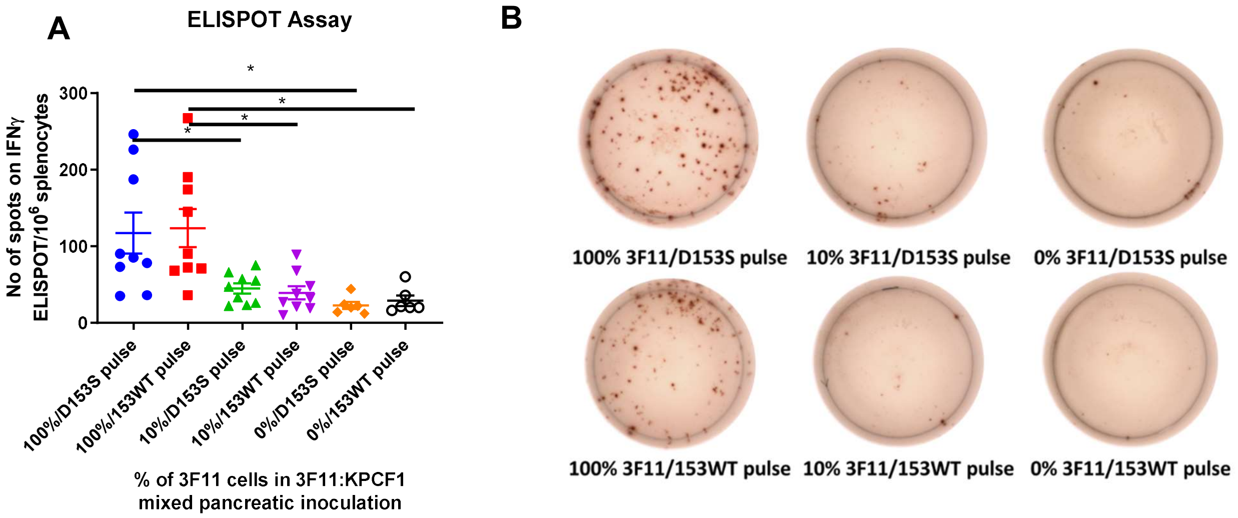


**Supplementary Fig. 1. Breaking tolerance to nontarget self-antigens at high frequency of neoantigens**, a) Results of Interferon Gamma ELISPOT on Day 10 post tumor inoculation. KPCF1 is a KPC cell line with $KRAS^{G12D}$ mutation, 3F11 is a KPC cell line with $KRAS^{G12D/D153S}$ mutation. Animals were sacrificed on Day 10 post orthotopic tumor inoculation, spleens were harvested (n = 3), and ELISPOT assay was performed. The splenocytes were pulsed with 153WT (VDDAFYTL) or D153S (VSDAFYTL) peptide. % denotes percentage of 3F11 cells, and pulse denotes peptide the splenocytes were pulsed with. For example, 100%/D153S pulse denotes splenocytes from mice inoculated orthotopically with 100% of 3F11 cells, pulsed with D153S peptide. Plates were read using AID ELISPOT reader, and quantified, b) Photographs of representative wells from ELISPOT plate, for each group. Data show mean ± SEM. * $p<0.05$.
